# Mathematical Analysis and Topology of SARS-CoV-2, Bonding with Cells and Unbonding

**DOI:** 10.1101/2021.05.07.443177

**Authors:** Arni S.R. Srinivasa Rao, Steven G. Krantz

## Abstract

We consider the structure of the novel coronavirus (SARS-Cov-2) in terms of the number of spikes that are critical in bonding with the cells in the host. Bonding formation is considered for selection criteria with and without any treatments. Functional mappings from the discrete space of spikes and cells and their analysis are performed. We found that careful mathematical constructions help in understanding the treatment impacts, and the role of vaccines within a host. Smale’s famous 2-D horseshoe examples inspired us to create 3-D visualizations and understand the topological diffusion of spikes from one human organ to another organ. The pharma industry will benefit from such an analysis for designing efficient treatment and vaccine strategies.

## 1. Introduction

The structure of the virus and spikes of the novel coronavirus (SARS-CoV-2 or COVID-19) that caused the suffering during 2020-2021 is understood in this article through topological constructions. We showed how such careful visualizations help to understand the virus-cell bonding through the distribution of spikes of the SARS-CoV-2. In general, the number of spikes and distribution of the spikes across various virus particles is found to be key in the spread of SARS-CoV-2 [1, 2], bonding of the spikes [3, 4, 5], and in understanding the entry of the virus into key organs like the lungs [7, 8, 9, 10]. We found that such a detailed mathematical analysis will eventually assist in the careful design of vaccines and medicines. Several studies analyze the situations of inactivity of the SARS-CoV-2 and the role played by the spikes [11, 12, 13, 14, 15, 16].

Many experimental results on the spikes and their activation, bonding, and inactivation assisted in vaccine development [17, 18, 19]. The pharmaceutical and the vaccine industry would benefit from such detailed visualizations of internal structures and bonding, rate of unbonding, and the role of interventions [20, 21, 22, 23].

In spite of experimental success in identifying a set of vaccine candidates for SARS-CoV-2 and the activity of the spikes, there exist several uncertainties in measuring successful vaccine impact. Theoretically, if a spike is completely bonded by an infected cell and this bonding is executed perfectly then that should lead to a new virus. At the same time, preventing a successful bonding and breaking of the spike would leave the virus incapable of spreading. Experiments leading to the identification of spike structures and their activities need to be more accurate and our present theoretical analysis promises to be useful for assisting in the experiments. Pharmaceutical and vaccination industries need to conduct highly accurate laboratory experiments. These experiments would need to carefully understand the role played by the spikes in SARS-CoV-2. Vaccines are designed to destroy the bonding capacity of these spikes or even destroy the spikes. Mathematical mapping, identification, and analysis of the spikes responsible for virus production within a host that are analyzed in this article are highly insightful for such experiments.

Our article will assist in improved design of vaccine experiments and better treatment designs that can take care of all the spikes at the time of entry into a host. Topological analysis is a rich tool and proper usage of it can help in avoiding uncertainties.

In this article, we have considered the following four assumptions in our topological constructions of the SARS-CoV-2:

i. Not all virus particles in the host are participating in infecting cells;
ii. Not all the spikes in a single virus may be bonded with cells;
iii. Each spike within a virus will bond with one and only cell;
iv. An empty spike (uninfected spike) of a given virus particle can bond with another cell.

This work provides original applications of topology which is one of the powerful tools of mathematical analysis [24, 25, 26, 27]. We cite here general references for the basic ideas of point-set topology. But our discrete constructions in this article are not explained in those sources. In the next section, we have described the basic topological space that we define using the number of spikes per virus particle within a host. We have provided novel usage of the mathematical analysis principles and topological constructions. A fraction of the spikes within a host in the space is allowed to get bonding with the uninfected cells. The entire structure of the space is mapped so that we can better understand the role of bonded and unbonded spikes within a host. Section 3 studies the role of treatment and vaccines in prohibiting the bonding and eliminating the infected host.

## 2. Topological Structures

Overall structure and topology of the virus and bonding/unbonding by cells within the host are described through Figure 2.2. Let 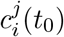 be the *i*^*th*^ novel coronavirus (ç) particle within a host at time *t*_0_ that has *j* number of spikes. We choose *i* = 1, 2, …, *n* and *j* = 1, 2, …, *j*_*i*_. Each of the spikes within the host is uniquely identified by this structure because 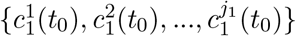 is the distinct set of spikes of the first virus particle and so on. In general, 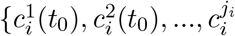 is the distinct set of spikes of the *i*^th^-virus particle for *i* = 1, 2, …, *n*.. Such a construction also allows us to write the expression:

**Figure 2.1.**
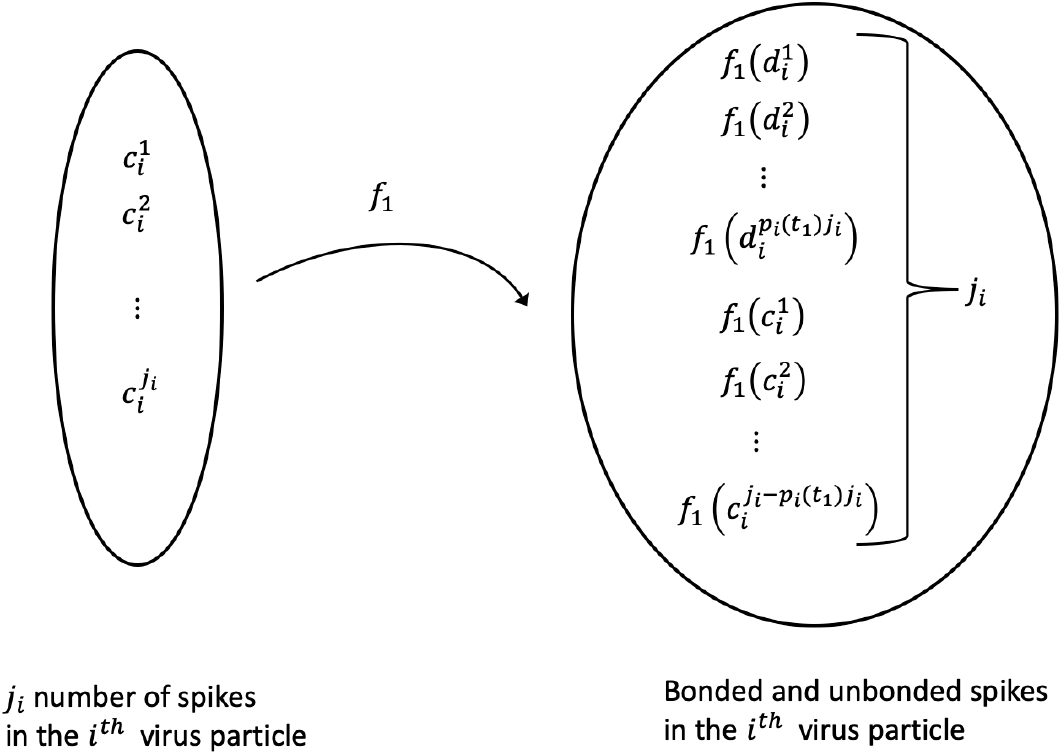
Mapping of spikes in the *i*^*th*^ virus particle at time *t*_0_ to bonded and unbonded spikes at time *t*_1_.

**Figure 2.2.**
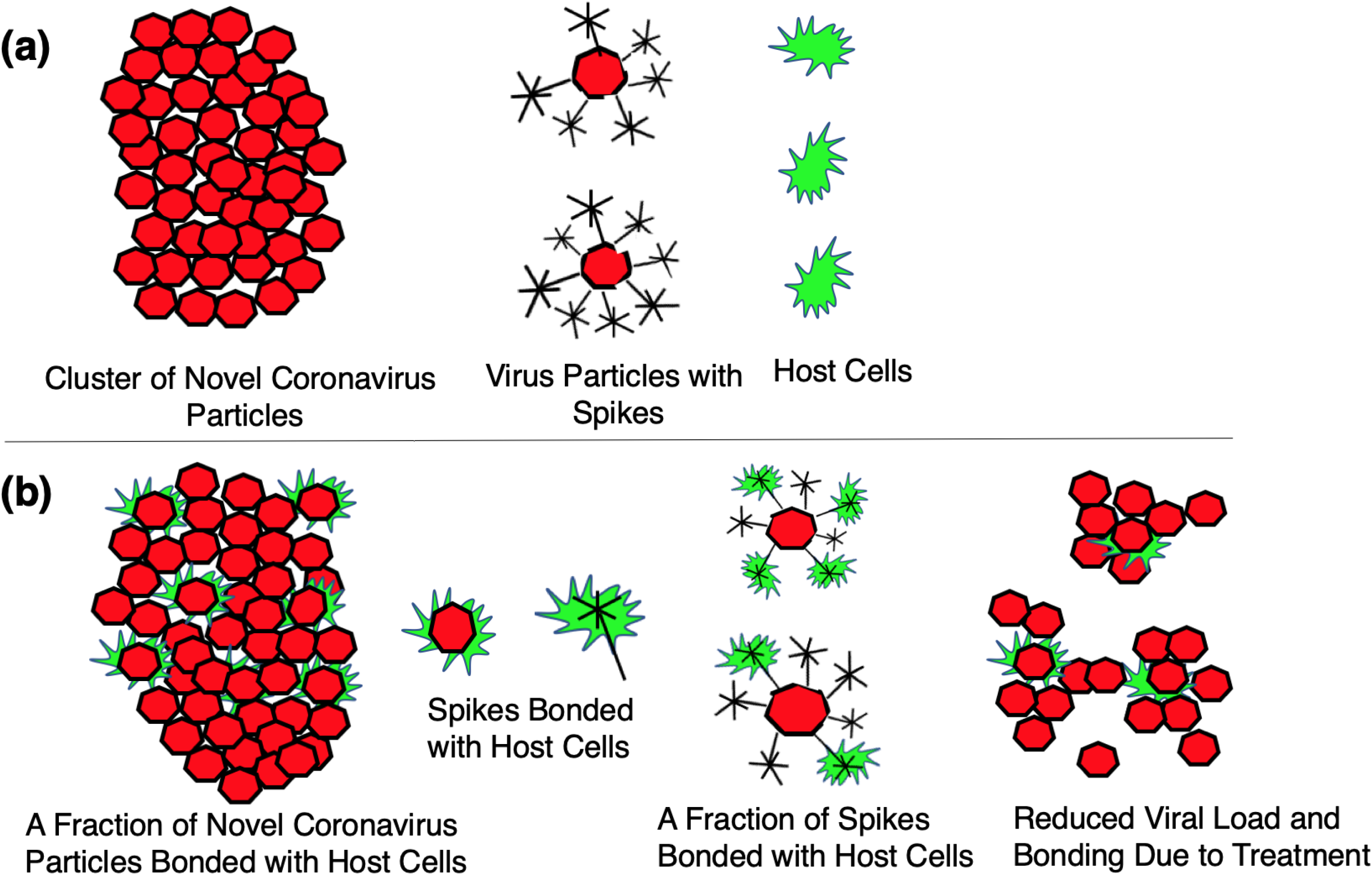
Topology of novel coronavirus (SARS-CoV-2), spikes, bonding, reduction in virus bonding due to treatment. (a) Imaginative description of the SARS-CoV-2, spikes and host cells, (b) Bonding of spikes to host cells, reduction of viral load due to treatment.

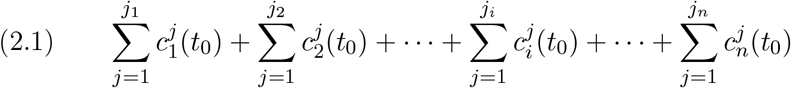

The quantity 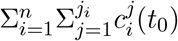 in (2.1) represents the total spikes in the host which are ready to bond with cells within the host. The spikes in the expression (2.1) are not yet bonded. Let *S*(*t*_0_) be the collection of all the spikes which were not yet bonded at time *t*_0_. Then

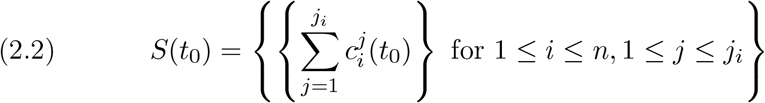

Let *p*_*i*_ be the fraction of the spikes in the *i*^th^-virus particle which are bonded with uninfected cells at time *t*_1_. The quantity *p*_*i*_ = 1 indicates that the *i*^th^ virus particle is fully bonded with uninfected cells, and each of the *j*_*i*_ spikes is occupied in bonding. The quantity *p*_*i*_ < 1 indicates that some of the spikes out of *j*_*i*_ spikes in the virus particle are unoccupied (or empty). We write

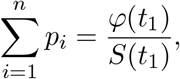

where *φ*(*t*_1_) is the total number of spikes in *S*(*t*_1_) which are bonded with uninfected cells at time *t*_1_. The cardinality of the set *S*(*t*_1_) is

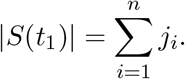

When *p*_*i*_(*t*_1_) = 1 for all *i*, then *φ*(*t*_1_) = |*S*(*t*_1_)| and hen *p*_*i*_(*t*_1_) < 1 for at least one *i*, then *φ*(*t*_1_) < |*S*(*t*_1_)|. Suppose that *f*_1_ : *S*(*t*_0_) *→ S*(*t*_1_), where *S*(*t*_1_) consists of the set of all spikes both bonded and unbonded. The number of spikes that were bonded during [*t*_0_, *t*_1_] is *p*_*i*_(*t*_1_)*j*_*i*_ and *j*_*i*_ *− p*_*i*_(*t*_1_)*j*_*i*_ is the number of spikes at *t*_1_ which are not bonded with uninfected cells for *i* = 1, 2, …, *n*. We assume an occupied spike with an uninfected cell will not be available for further bonding. So the bonded spikes at *t*_1_, i.e., *p*_*i*_(*t*_1_)*j*_*i*_, have completed their virus bonding capacity by time *t*_1_, and the remaining spikes available at time *t*_1_ are *j*_*i*_ *− p*_*i*_(*t*_1_)*j*_*i*_. These unbonded spikes will be available for bonding during (*t*_1_, *t*_2_]. Let 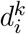 be the *k*^*th*^ bonded spike at time *t*_1_ out of *j*_*i*_ spikes at time *t*_0_ for *k* = 1, 2, …, *p*_*i*_(*t*_1_)*j*_*i*_ such that

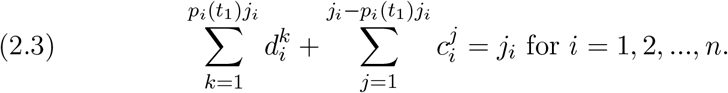

The number of occupied spikes among all the virus particles during [*t*_0_, *t*_1_] which will not be available for further bonding are

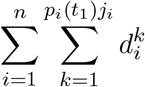

If *p*_*i*_(*t*_1_) = 1 for any *i* at time *t*_1_ then that virus particle has completed the bonding role in the system. It is assumed that *p*_*i*_(*t*_1_)*j*_*i*_ number of spikes for each *i* (when *p*_*i*_(*t*_1_) < 1) would generate *p*_*i*_(*t*_1_)*j*_*i*_ number of new virus particles available for bonding during [*t*_0_, *t*_1_], else (when *p*_*i*_(*t*_1_) = 1) it would generate *j*_*i*_ number of new virus particles during the same period. See Figure 2.1.

Hence the newer virus particles produced during [*t*_0_, *t*_1_] are 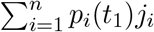,so the available virus particles for bonding at time *t*_1_ will be

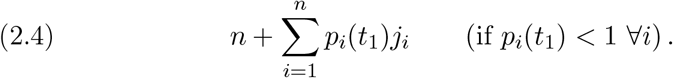

The birth rate *λ*_*i*_(*t*_1_) (*w*.*r*.*t. n*) of the new virus particles during [*t*_0_, *t*_1_] is

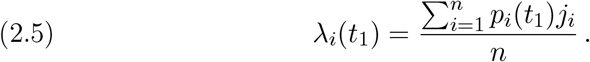

Not all of the new viruses at time *t*_1_ in (2.4) may be available for bonding during (*t*_1_, *t*_2_] if one or more of the virus particles (out of *n*) might have all its spikes bonded at time *t*_1_.

Suppose that *p*_*i′*_ (*t*_1_)*j*_*i′*_ = *j*_*i′*_ for *i*′ = 1, 2, …, *m* (*m* < *n*) and *p*_*i\**_(*t*_1_)*j*_*i\**_ < *j*_*i\**_for *i\** = 1, 2, …, *n − m*, such that

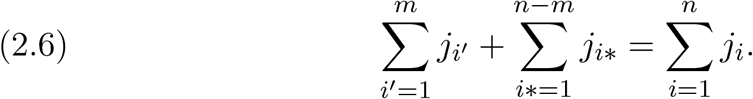

Based on (2.6), the number of virus particles available at time *t*_1_ after removing completely bonded virus particles during [*t*_0_, *t*_1_] (i.e., those virus particles for which all of its spikes were bonded with uninfected cells, and adding new virus particles created by the *n−m* virus particles with available spikes for bonding during [*t*_0_, *t*_1_]) are

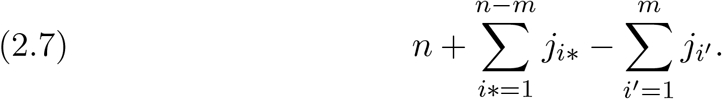

The total number of bonded spikes due to 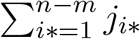 and 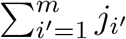 are

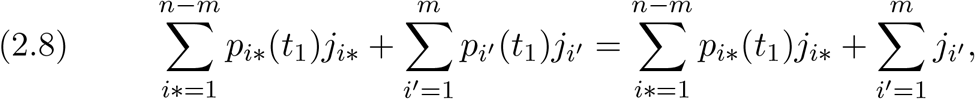

and, from (2.3),

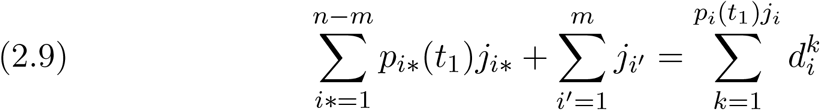

Let us decompose 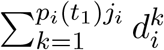 as

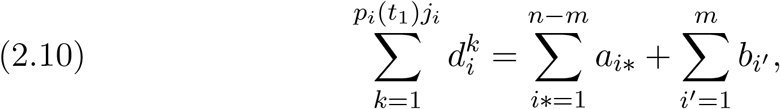

where *a*_*i\**_ = *p*_*i\**_(*t*_1_)*j*_*i\**_ and *b*_*i′*_ = *j*_*i′*_.

The total bonded during [*t*_0_, *t*_2_] is responsible for giving birth to new viruses as described previously. The total number of remaining spikes, those |*S*(*t*_1_)| which are available at time *t*_1_, are the sum of (i) the number of spikes unbonded during [*t*_0_, *t*_1_], and (ii) the number of spikes that are created due to the birth of new virus particles. The listing of the set *S*(*t*_1_) of spikes helps in constructing the function *f*_2_ : *S*(*t*_1_) *→ S*(*t*_2_). Here *S*(*t*_2_) is the set of spikes created by *S*(*t*_1_) during (*t*_1_, *t*_2_]. Let us list the elements of the set *S*(*t*_1_) below:

The list of unbounded spikes *J*_1_ during [*t*_0_, *t*_1_] is obtained as remaining spikes from the first term of (2.9) as

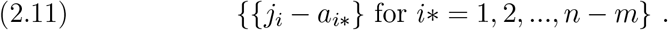

The list of spikes *A*_1_ available at *S*(*t*_1_) because of the first term of the R.H.S. of (2.10) is

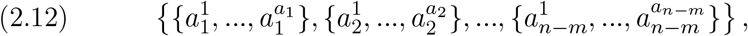

Where the 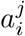 in (2.12.) represent the *j*^*th*^ spike of the *i*^*th*^ virus resulting from 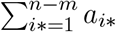 in (2.10). That is, as per the set in (2.12), there are number of spikes for the first virus, *a*_2_ numbr of spikes for the second virus, and so on *a*_*n−m*_spikes for the (*n − m*)^*th*^ virus. The list of spikes *B*_1_ available at *S*(*t*_1_) due to the resultant of the second term of the R.H.S. of (2.10) is

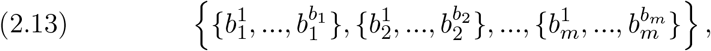

where the 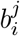 in (2.13) represent the *j*^*th*^ spike of *i*^*th*^ virus resulting out of 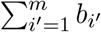 in (2.10). That is, as per the set in (2.13), there are *b*_1_ number of spikes for the first virus, *b*_2_ numbr of spikes for the second virus, and so on *b*_*m*_ spikes for the *m*^*th*^ virus. The domain *S*(*t*_1_) of the function *f*_2_ will have the collection of all the elements of the sets *J*_1_, *A*_1_ and *B*_1_, i.e.

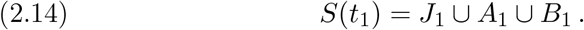

The collections *S*(*t*_0_) and *S*(*t*_1_) constructed above can be treated as two spaces and *S*(*t*_1_) in (2.14) is now seen as a disconnected space.

### Lemma 1

*The function f*_1_ *is not* 1*–*1 *when bonding occurs during* [*t*_0_, *t*_1_].

*Proof:* We have *f*_1_ : *S*(*t*_0_) *→ S*(*t*_1_), where *S*(*t*_0_) and *S*(*t*_1_) are the sets of all the distinct spikes at time *t*_0_ and time *t*_1_. When bonding occurs during [*t*_0_, *t*_1_], the set of all spikes at time *t*_1_ will be *S*(*t*_1_) as seen in (2.14). This implies that

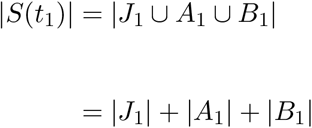

From (2.11) to (2.13) we can write

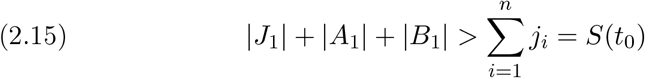

Because of the inequality (2.15), *f*_1_ cannot be 1–1.

□

### Corollary 2

*The function f*_1_ *is* 1*–*1 *when no bonding occurs during* [*t*_0_, *t*_1_]. *When the bonding does not occur then* |*S*(*t*_0_)| = |*S*(*t*_1_)| *and the elements of S*(*t*_0_) *and S*(*t*_1_) *are not different*.

Since we can consider

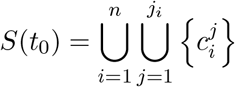

and the elements of *s*(*t*_0_) are distinct, we treat here *S*(*t*_0_) as a discrete topological space with 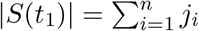 elements in the space *S*(*t*_0_). Let *S*_*X*_(*t*_0_) and *S*_*Y*_ (*t*_1_) be two subsets of *S*(*t*_0_) such that *S*_*X*_(*t*_0_) represents bonded spikes and *S*_*Y*_ (*t*_0_) represents unbonded spikes during [*t*_0_, *t*_1_]. Then, by the construction of *S*(*t*_0_), the two subsets *S*_*X*_(*t*_0_) and *S*_*Y*_ (*t*_0_) form two disjoint subspaces of *S*(*t*_0_). The space *S*(*t*_1_) as well is a discrete topological space and three subsets of it *J*_1_, *A*_1_ and *B*_1_, form three disjoint topological discrete subspaces of *S*(*t*_1_).

### Definition 3. Topological diffusion

We define here the topological diffusion *D*_*s*_(*t*_0_) of the space created due to newer spikes during [*t*_0_, *t*_1_], as

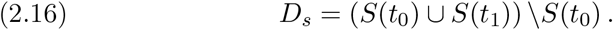

**Theorem 4**

*The topological diffusion D*_*s*_(*t*_0_) *during* [*t*_0_, *t*_1_] *is A*_1_ ∪ *B*_1_.

*Proof*

We have

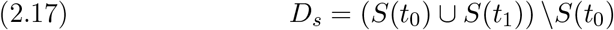

The set *D*_*s*_(*t*_0_) indicates the newer elements created in the combined space (*S*(*t*_0_) ∪ *S*(*t*_1_)). The collection of elements of *S*(*t*_0_) and *S*(*t*_1_) in (2.17) are further expressed using (2.14) as follows:

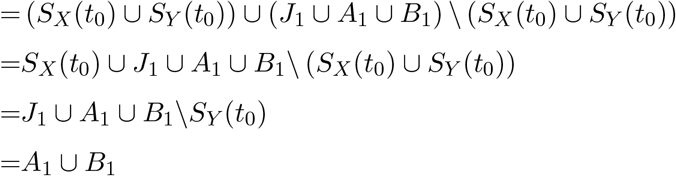

□

The collection *A*_1_ ∪ *B*_1_ is the newer space created during [*t*_0_, *t*_1_].

### Example 5

Suppose *S*(*t*_0_) has 10 spikes. The topological diffusion occured during [*t*_0_, *t*_1_] to arrive at |*S*(*t*_1_)| = 21. See Figure 2.3 for mapping of bonded cells into new spikes and carrying forward the unbonded spikes.

**Figure 2.3.**
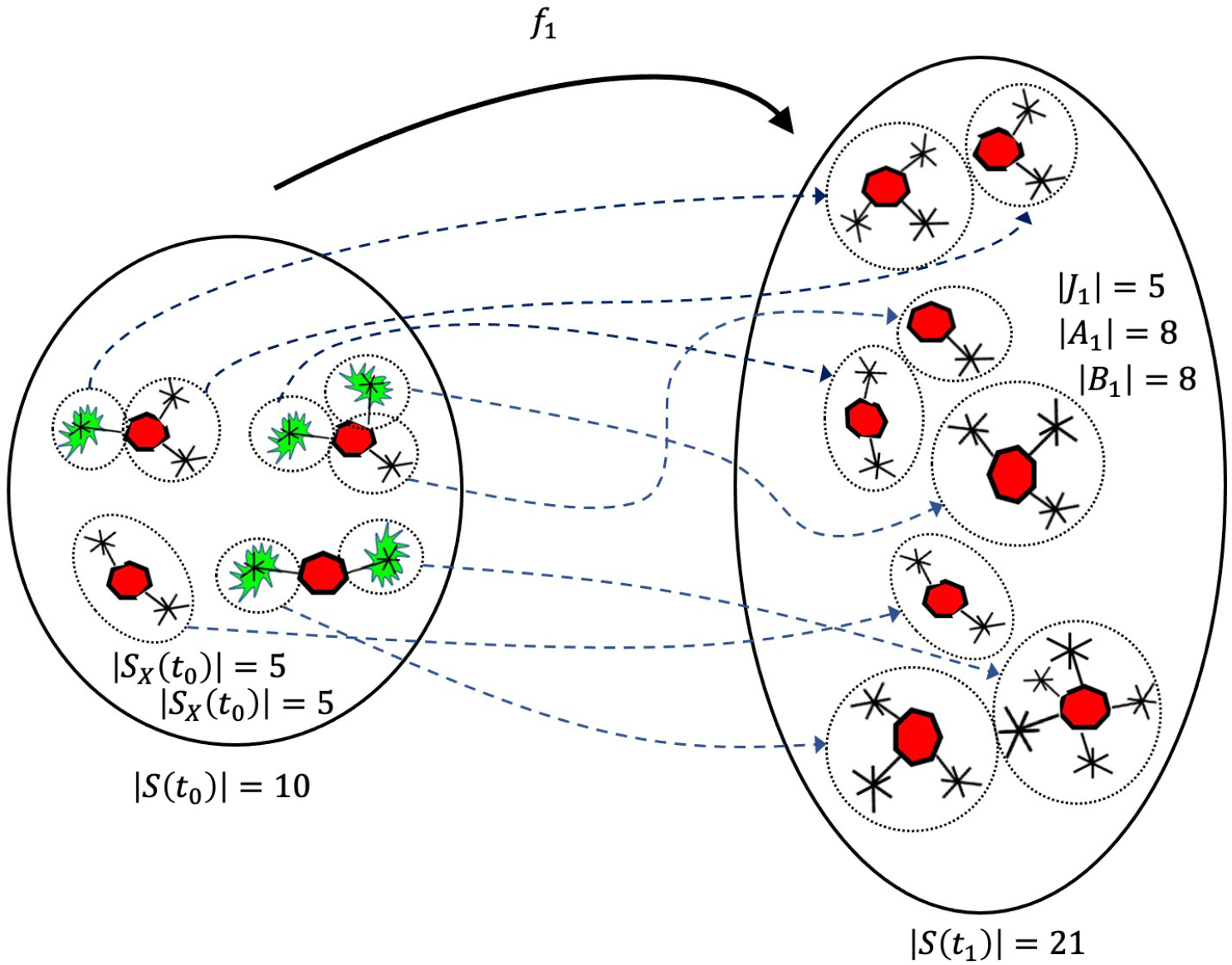
Numerical example of mapping of bonded and unbonded spikes. *A*_1_ ∪ *B*_1_ = 16 as shown is the topological diffusion of the space.

Every singleton set within *S*(*t*_0_) is an open subset. That means that each spike in *S*(*t*_0_) is considered as a singleton set and *S*_*X*_(*t*_0_) and *S*_*Y*_ (*t*_0_) form two open subsets of *S*(*t*_0_). In fact, according to discrete topology *S*_*X*_(*t*_0_) and *S*_*Y*_ (*t*_0_) can also be treated as closed subsets (so the space is disconnected). The transformations of the space *S*(*t*_0_) during [*t*_0_, *t*_1_] would lead to newer spaces due to bonding (also argued as in the proof of Lemma 1). Such a creation of new topological spaces and their cardinality can be influenced with a treatment intervention at some time *t* for *t* ∈ [*t*_0_, *t*_1_]. Treatment works in reducing the value of *p*_*i*_ or the death rates of the virus particles 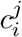 or both.

## 3. Treatment and Vaccinations

We assume a treatment to kill the virus population (viral load within a host) would increase the mortality rate of the virus population and reduce the bonding of the uninfected cell population with SARS-CoV-2. At time *t*_0_ the viral load would be lower and treatment during [*t*_0_, *t*_1_] would have a higher impact on reducing the viral load than if the treatment was introduced during [*t*_1_, *t*_2_]. We assume a longer time to introduce a treatment after time *t*_0_ according to the longer time the virus population is restricting the virus growth. Since |*S*(*t*_0_)| < |*S*(*t*_1_)| in the absence of treatment, *S*(*t*_0_) = *S*(*t*_1_) can be achieved (Corollary 2) when treatments are introduced during [*t*_1_, *t*_2_]. We assume the host will be dominated by the virus when virus growth is not controlled and the virus will be eliminated either naturally or due to treatment impact. Let *p*_*i*_ be defined as earlier and let *q*_*i*_(*t*_1_) be the fraction of the spikes in the *i*^*th*^ virus which are bonded during [*t*_0_, *t*_1_] and treatment was introduced at some time *s* for *s ∈* (*t*_0_, *t*_1_]. Here, 0 ≤ *q*_*i*_(*t*_1_) < *p*_*i*_(*t*_1_). The quantity *q*_*i*_(*s*) = 0 means there are no bonded spikes at *s*. The quantity *q*_*i*_(*s*) would never reach *p*_*i*_(*t*_1_). Since the treatment would also increase the mortality rate of the 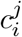 population, we assume that *q*_*i*_(*t*_1_) = 0 if *i*^th^ virus dies at *s* for *s* ∈ (*t*_0_, *t*_1_] or *q*_*i*_(*t*_1_) = 0 if no bonding with the spikes of the *i*^*th*^ virus occurs. When *q*_*i*_(*t*_1_) = 0, then all the spikes of the *i*^*th*^ virus at *t*_0_ will be available for bonding during (*t*_0_, *t*_1_].

### Theorem 6

*Consider the sequence of functions* (*f*_*n*_)_*n*≥1_, *where f*_*n*_ : *S*(*t*_*n−*1_) *→ S*(*t*_*n*_). *Suppose a treatment is introduced at t*_*m*_ *for some m* ≥ 1. *Then the sizes of S*(*t*_*i*_) *are increasing until t*_*m*_ *and the sizes of S*(*t*_*i*_) *are decreasing until S*(*t*_*m−*1_) *and f*_*m*_ : *S*(*t*_*m−*1_) *→ φ (empty set)*.

*Proof*. When treatment prohibits bonding then virus population having no host would eventually die. Treatment would also kill bonded cells with spikes. Suppose the treatment is introduced at time *s* for *s* > *t*_0_ and

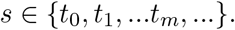

The time intervals were partitioned as {[*t*_0_, *t*_1_], (*t*_1_, *t*_2_], …,}. Given that *s* = *t*_*m*_, we assume impact of the treatment can be measured in prohibiting the bonding at *t*_*m*_ for *m >* 0. Similarly the treatment would have impact on killing the bonded cells with spikes at *t*_*m*_. Hence

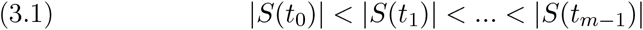

and

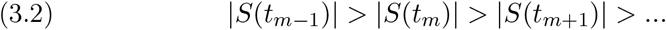

The sequence *S*(*t*_*m−*1_)_*m*≥1_ in (3.2) is monotonic and 0 ≤ |*S*(*t*_*m*_)| for all *m* ≥ 0. Hence, by the monotone convergence theorem, the sequence (3.2) is convergent to *φ* (empty set).

### Theorem 7

*Consider the sequence of functions* (*f*_*n*_)_*n*≥1_, *where f*_*n*_ : *S*(*t*_*n−*1_) *→ S*(*t*_*n*_). *Suppose the host is vaccinated prior to t*_0_. *Then the sequence S*(*t*_*m*_)_*m*≥0_*is decreasing and lim*_*n*→∞_ |*S*(*t*_*m*_)| = 0.

*Proof*. Given *f*_*n*_ : *S*(*t*_*n−*1_) *→ S*(*t*_*n*_) for all *n*. When the host is vaccinated prior to *t*_0_ the system will prohibit the spikes to get bonded with cells. Unbounded spikes and virus particles dying over time lead to the decreasing sequence

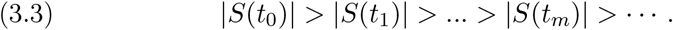

Similar to the argument of the proof of the Theorem 6, we have

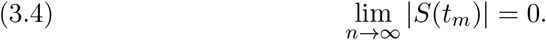

### *Remark* 8

A vaccinated host would alter the range of *f*_1_ in Example 5 but the domain of *f*_1_ consisting of vaccinated and not vaccinated hosts remains the same. COVID-19 vaccinations would not prevent virus to enter an unprotected host, but a vaccinated host prevents the virus from getting bonded with the virus spikes.

Given Theorem 7, this sequence of inequalities will emerge:

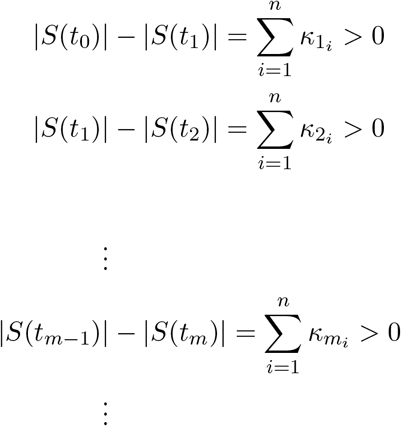

Since (3.4) is true, we will have

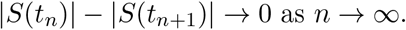

Here 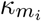 for *m* = 1, 2, … is the remaining number of spikes unbounded during [*t*_*i−*1_, *t*_*i*_] for *i* = 1, 2, …, *n*. We can create the elements similar to (2.11) to (2.13) for the periods {[*t*_0_, *t*_1_], (*t*_1_, *t*_2_], …,}. Let *J*_*τ*_, *A*_*τ*_, *B*_*τ*_ be the sets defined on the intervals {[*t*_0_, *t*_1_], (*t*_1_, *t*_2_], …,} similar to *J*_1_,*A*_1_,*B*_1_ which were defined from (2.11) to (2.13) for the emenets defined on the interval [*t*_0_, *t*_1_]. The topological diffusion created until the treatment initiated at *t*_*m*_ is 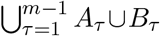. Hence the diffusion of the elements created will start declining with the initiation of the treatment. The smaller the value of *t*_*m*_, the lesser the quantity 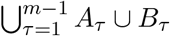.

### Theorem 9

*Topological structures of the virus populations and spikes will be different under vaccination and treatment of hosts even though* |*S*(*t*_*n*_)| *−* |*S*(*t*_*n*+1_)| *→* 0 *as n →* ∞.

*Proof*. Suppose the treatment is initiated within a host at *t*_*m*_ for *m* ≥ 1. The topological structure of spikes would first have the increasing property in (3.1), and then will start decreasing as in (3.2). This leads to |*S*(*t*_*n*_)| *−* |*S*(*t*_*n*+1_)| *→* 0 as *n →* ∞.

Under a vaccinated host, as soon as the SARS-CoV-2 virus enters at *t*_0_, the topological structure of the spike population spread within the host obeys (3.3), and that leads to |*S*(*t*_*n*_)| *−* |*S*(*t*_*n*+1_)| *→* 0 as *n →* ∞.

Hence, two topological structure described above will be different although the limiting number of spikes diminishes.

### 3.1. Horseshoe mapping

Inspired by Stephen Smale’s original famous *horseshoe* example [28, 30, 31], we have visualized a discretized version of the same idea with plastic beads in a container. Let us consider a hollow cube and fill it with plastic beads. Suppose all the beads in this cube are transferred into a horseshoe-shaped pipe. See Figure 3.1. Note that the original horseshoe mapping is continuous and is a diffeomorphism between a square and horseshoe-shaped space. Let us imagine the size of the *S*(*t*_1_) SARS-CoV-2 spikes are located in the throat area of a human host. Suppose during the interval (*t*_1_, *t*_2_] these spikes are spread into the lung area. Assume that the treatment to control the virus is initiated at *t*_2_ such that the throat area spikes are eliminated during (*t*_2_, *t*_3_] and the number of spikes at *t*_3_, is the set *S*(*t*_3_). This leads to

**Figure 3.1.**
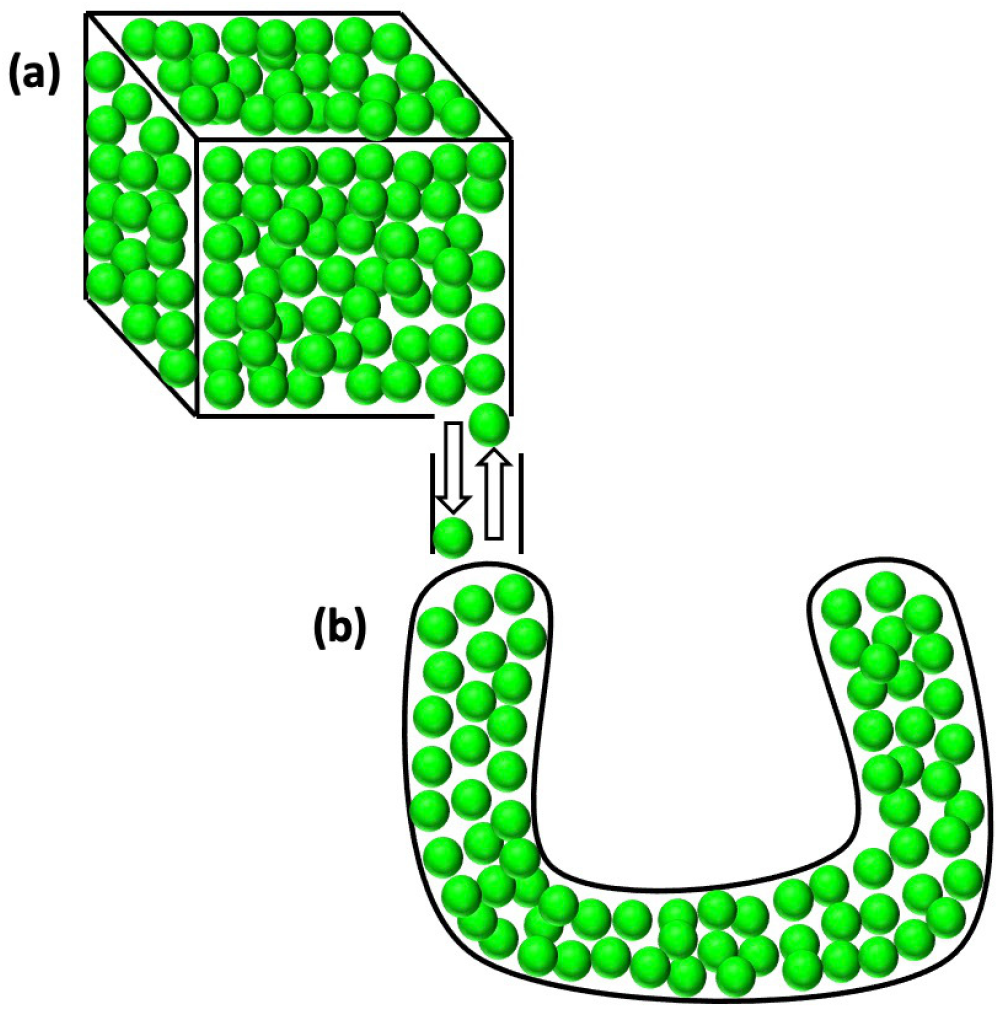
Transformation of a cube containing beads into a horseshoe-shaped pipe containing the same number of beads. **(a)** A cube container of green color beads, **(b)** Beads in **(a)** are transferred into a horseshoe shaped pipe. The number of beads in **(a)** and **(b)** are equal. The beads in **(b)** can be transferred into **(a)**. The beads occupying capacities of containers in **(a)** and **(b)** are equal and it is assumed that no free space left to add an additional bead into these containers. The original example of Stephen Smale is a diffeomorphism between two 2-D objects, namely, a square and a horseshoe, famously known as Smale’s *horseshoe*.

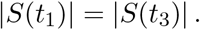

Transformation of the number of spikes of *S*(*t*_1_) in the throat area into number of spikes of lungs area is demonstrated in Figure 3.2. Suppose the size of the spikes at *t*_1_ are located in the throat area of a host. We have *f*_1_ : *S*(*t*_0_) *→ S*(*t*_1_). Under the no treatment assumption during (*t*_1_, *t*_2_], we have *f*_2_ : *S*(*t*_1_) *→ S*(*t*_2_), where |*S*(*t*_2_)| *>* |*S*(*t*_1_)|. The newer space created during (*t*_1_, *t*_2_] is

**Figure 3.2.**
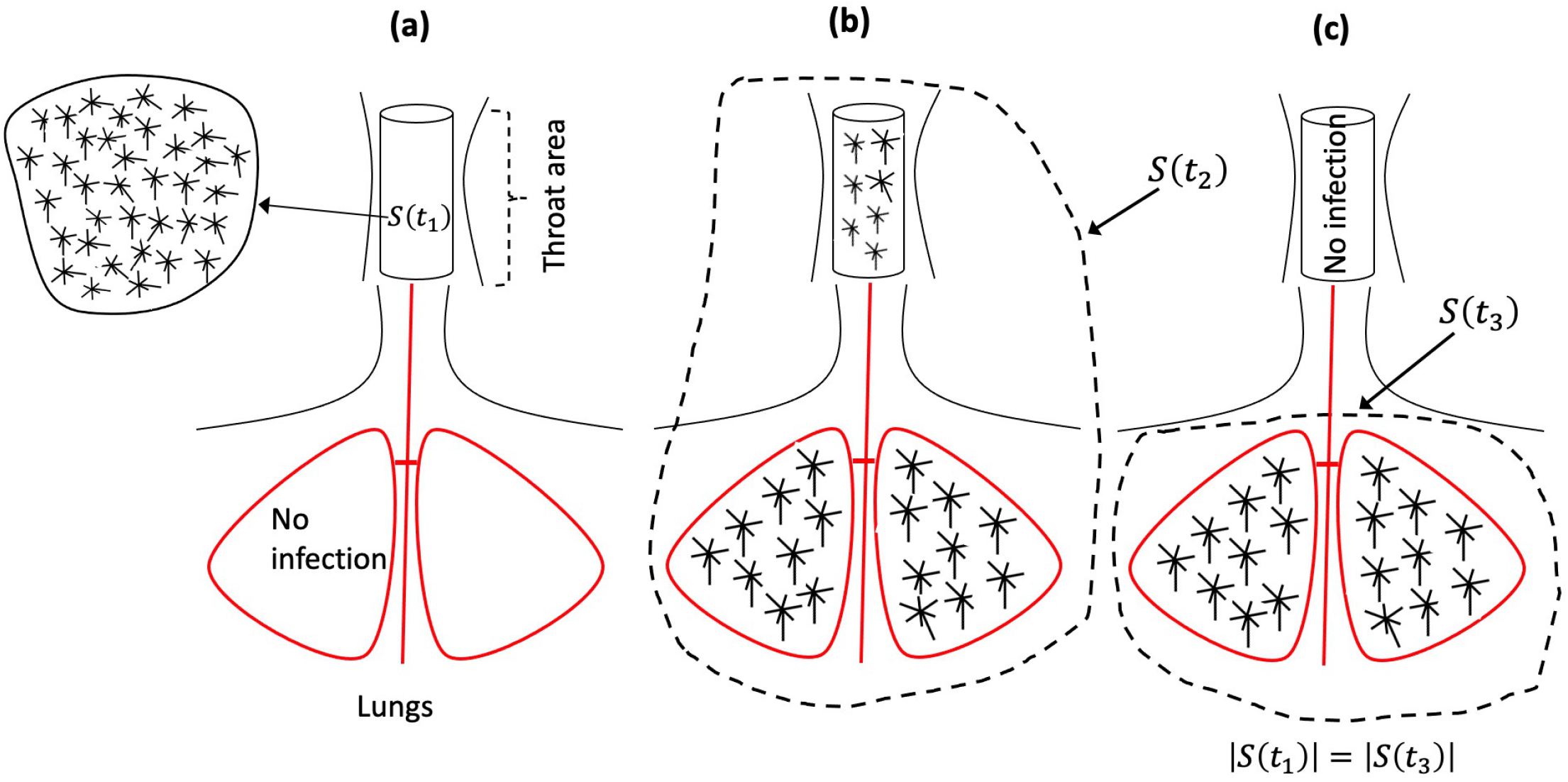
Transformation of the number of spikes at *t*_1_ in the throat area into the number of spikes at *t*_3_ in the lungs area. (*a*) The number of spikes in the throat at *t*_1_ are *S*(*t*_1_) and no infection in lungs, (*b*) The number of spikes grown during (*t*_1_, *t*_2_] expanded into lungs and treatment is introduced at *t*_2_, (*c*) The spikes in the throat are killed and the number of spikes at *t*_3_ in the lungs are *S*(*t*_3_) and |*S*(*t*_1_)| = |*S*(*t*_3_)|.

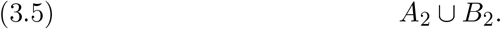

The topological diffusion in (3.5) gives us,

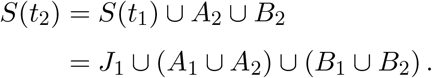

Suppose the treatment for SARS-CoV-2 is introduced at *t*_2_ such that

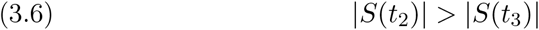

is attained. There are now three possibilities that will arise due to (3.6):

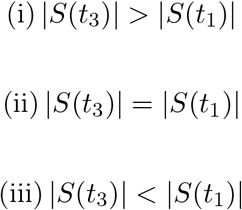

Above possibility (ii) we associate with that of Smale’s *horseshoe* type of example and is also demonstrated in Figure 3.2. During (*t*_2_, *t*_3_] the number of spikes killed in the throat due to the treatment initiated at *t*_2_ and due to the creation of the topological diffusion *A*_2_ ∪ *B*_2_ reaches the size |*S*(*t*_1_)| at *t*_3_. Then this kind of discrete topological transformation of the number of spikes at *t*_1_ into the number of spikes at *t*_3_ in a different location of a host is topologically visualized as an *horseshoe* type of example. Of course, we are aware in a true sense Smale’s *horseshoe* is a diffeomorphism between two open spaces (a square and a a horseshoe). The current analysis of transformations of the number of spikes located at *t*_1_ in the throat area and the number of spikes at *t*_3_ in the lungs within a host handles the points (elements) of the space discretely.

The horseshoe example transforms the points of an open square into an equivalent area horseshoe using continuous mapping. The implications of the horseshoe are plenty—for example, the squeezing and stretching of a square to a horseshoe-shaped space in the creation of hyperbolic dynamics and chaos. In our analogy, the spikes in the throat within a human host do not get transferred to the lungs because virologically spikes do not travel within the host but they grow over time under a no-treatment scenario. Only after treatment is initiated are the spikes in the throat killed and an equivalent number of newer spikes born to remain active for some time in the lungs. We imagine this phenomenon as described through Figure 3.2 as the transformation of spikes of the throat to that of the lungs.

The geometry of the horseshoe is especially meaningful for us because spikes from the throat area due to the initial infected virus population are all located in one place. Then, due to the spread of the virus over the intervals {[*t*_0_, *t*_1_], (*t*_1_, *t*_2_], …,}, they diffuse into different organs which have geometrically a different shape than the throat. With the example of beads (Figure 3.1), and assuming no scope for adding a new bead in the cube, the corresponding pipe would take the 2D horseshoe to a 3D similar-shaped pipe through discrete topology. Our spikes analogy is that the horseshoe example was built on 2D and diffusion of spikes within the human organs is imagined in 3D.

## 4. Discussion

Our study provides the most accurate mathematical structuring of the space of the SARS-CoV-2 virus and its spikes within the host. See the expression 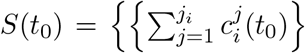 for 1 ≤ i ≤ *n*, 1 ≤ *j* ≤ *j*_i_ in section 2 and the clustering of spikes into bonded and unbonded spikes. The advantage of such a structure is to provide a detailed scope for mapping a spike that is available for bonding with an uninfected cell. Not a single unbonded spike will be left out in this process. The procedure also helps in tracking a bonded virus in such a way that the birth of newer virus particles through bonded spikes is monitored. See the expression *S*(*t*_1_) = *J*_1_ ∪ *A*_1_ ∪ *B*_1_, where *J*_1_, *A*_1_, and *B*_1_ form three disjoint topological discrete subspaces mapped out from *S*(*t*_0_). The set *J*_1_ emerges out of unbonded spikes in a previous time point and *A*_1_ ∪ *B*_1_ is the collection of spikes generated due to bonding spikes with cells at a previous time point. Our clear-cut visualization of the theoretical constructions helps in understanding the structure of the spike-cells within the space.

Laboratory experiments on the virus particles and bonding are usually done on a group of viruses. Our procedure provides deeper insight for better design in conducting experiments on isolated individual viruses. Such a method will help in aggregating the virus population along with their number of spikes and measuring bonded and unbonded spikes for each virus particle. Lemma 1 provides the growth of spikes and their mapping of initial spikes that can create newer spaces within a time interval.

One of the central features of our construction is the development of a new measure that we call “*topological diffusion*.” In general topology, no such measure exists. Using this measure, one can study the growth of spikes over time. The topological diffusion introduced in this article not only identifies the new spikes that emerge but how many of those were due to virus particles that had partial bonding of the spikes. These novel ideas make our work more practically implementable in pharmaceutical and vaccine industrial experiments. We have theoretically established this value within a small interval *A*_1_ ∪ *B*_1_ and also over a large interval. The descriptions of *A*_1_and *B*_1_ are recorded in previous paragraphs and also in section 2. Figure 2.3 provides an example of measuring the topological diffusion.

Theorem 6 provides the impact of a treatment in eliminating virus particles and Theorem 7 provides the impact of the vaccine on eliminating viruses after entry into a host. We also generalize our results over multiple time intervals and the timing of initiation of therapy. Topological diffusion constructed in the article is also associated with Stephen Smale’s famous horseshoe type of example. The original example by Smale was constructed as a diffeomorphism of two open spaces, namely, a square and corresponding sized area of a horseshoe. However, the current article considered a transformation of a discrete collection of spikes in one organ of a human host into the equivalent number of spikes in a different organ. Moreover, the demonstration we provided was between two 3-D shaped organs within a human host. Such visualization of the horseshoe example is new in the literature.

### Concluding Remarks

The study presented in this paper is original and incisive. It uses powerful mathematical techniques—most notably ideas from topology—to analyze the bonding of corona virus cells. Our emphasis on discrete topology is somewhat novel.

As a result we obtain insights that will be useful in the production of new and more effective vaccines. We believe that the use of mathematical analysis in a medical context is a new and effective technique for epidemiology that will become recognized and solidly established in future work.

